# Estimating the overlap between two malaria parasites’ *var* repertoires

**DOI:** 10.1101/343152

**Authors:** Daniel B. Larremore

**Affiliations:** Department of Computer Science, University of Colorado at Boulder; BioFrontiers Institute, University of Colorado at Boulder

## Abstract

Measuring the overlap between the *var* gene repertoires of two *P. falciparum* parasites is, in principle, easy. Each parasite genome contains a repertoire of approximately 60 *var* genes, so upon fully sequencing both parasites’ genomes, the number of shared *var* sequences can be directly counted. In practice, however, only a fraction of each parasite’s *var* repertoire is likely to be sampled due to the difficulties of whole-genome sequencing for *var* genes and the stochastic sample provided by PCR techniques. Although a method exists for quantifying repertoire overlap under these subsampled conditions, its bias is well documented and the uncertainty of its estimates cannot be quantified. Here we derive and validate a method to rigorously estimate the repertoire overlap between two parasites from the overlap of their subsampled repertoires. By solving a Bayesian inference problem, this method takes into account the rates of subsampling and produces unbiased and Bayes-optimal estimates of overlap. In addition, it provides a natural framework for computing the uncertainty of its estimates, and can be used in laboratory planning by quantifying the tradeoff between sequencing effort and uncertainty.

## I. INTRODUCTION

Of the diverse multigene families of *Plasmodium falciparum*, the *var* gene family is the most heavily studied because of its direct links to both malaria’s duration of infection and its virulence. When a *var* gene is expressed, the protein product is exported to the surface of the infected red blood cell where it facilitates binding to endothelial cells, allowing the parasite to sequester itself away from free circulation. Furthermore, each parasite’s ~ 60 *var* genes are hypervariable and mutually distinct, allowing parasites to epigenetically switch expression from one *var* gene to another when the former begins to elicit a strong immune response [1, 2]. While these immune evasion strategies are key to prolonging an untreated infection, *var* genes have also been linked to malaria’s most deadly symptoms. In particular, because individual *var* gene products bind with varying affinity to tissuespecific endothelial cells [3, 4], expression levels of different subtypes of *var* genes are linked to malaria’s worst clinical phenotypes like coma, anemia, and respiratory distress [5, 6].

Recent studies of *P. falciparum* epidemiology and evolution have begun to learn from comparisons of the sets of genomic *var* repertoires between parasites [7–13]. Since *var* repertoires are, themselves, under selection, theory suggests that if a human population has been exposed to particular *var* genes, then repertoires that contain those *var* genes will have a lower fitness than repertoires that are entirely unrecognized by local hosts, shaping the *var* population structure [14, 15]. In fact, preliminary studies have found evidence to support the idea that population immunity is shaping the content of *var* repertoires [10, 11], paving the way for these hypervariable gene repertoires to be used in high-sensitivity genetic epidemiological studies. A followup study to Ref. [10] found that *var* repertoires in a region of decreasing transmission have coalesced into highly related clusters that appear to be non-overlapping with each other, driven by local immune adaptation [13], although at the time of writing this study remains under peer review (June, 2018). Methods by which we estimate the extent to which *var* repertoires overlap are therefore important, particularly as studies of the population genetics and genetic epidemiology of malaria’s antigens become more sophisticated and data rich.

Measuring the overlap between repertoires would be straightforward if it were easy to fully sequence a parasite’s *var* genes. However, due to their massive diversity and recombinant structure, assembly of *var* genes is difficult—indeed, whole-genome sequencing’ for *P. falciparum* often excludes *var* genes because they are simply too difficult to assemble. Instead, experimentalists use degenerate PCR primers targeting a small “tag” sequence within a particular domain called DBL*α* [16] which is found in almost all *var* genes. Due to their experimental accessibility, DBL*α* tags have been widely used to study the structure and function of *var* genes [1, 6, 7, 10, 13, 16–19]. Still, these PCR techniques generate a random sample of 60 or fewer unique tag sequences from each parasite. This means that experimental measurements of repertoire overlap are performed using stochastic subsamples whose *empirical* overlap may fluctuate from experiment to experiment (Figure 1), motivating the three aims of this paper. First, how can we estimate the true overlap between repertoires when we can only measure the overlap between samples from repertoires? Second, how can we quantify the uncertainty around our repertoire overlap estimates? Third, what are the implications of uncertainty for the design and budgeting of *var* tag studies?

**FIG. 1.**
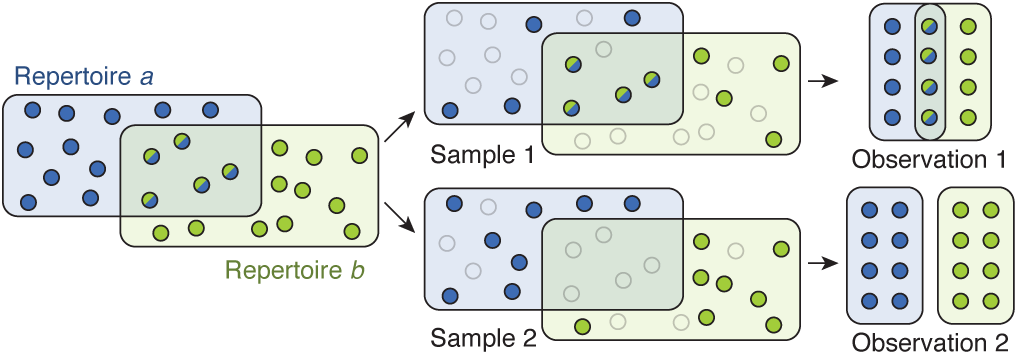
Stochastic sampling leads to variation in observed repertoire overlap. The genes of two hypothetical repertoires are represented by blue and green circles, respectively. Each repertoire has 16 genes, and *s* = 5 genes are members of both repertoires. In two independent sampling experiments, shown in top and bottom rows, *n_a_* = *n_b_* = 8 genes are sampled at random from each repertoire (dark circles) while the other 8 genes are not sampled (transparent circles). Observation of the first experiment finds a repertoire overlap of *n_ab_* = 4, while observation of the second finds *n_ab_* = 0.

The only existing method to compute repertoire overlap, called *pairwise type sharing* [7] in the malaria literature, is extremely intuitive. Suppose that PCR methods have produced *n_a_* and *n_b_* tags from parasites *a* and *b*, respectively, and that a sequence-level comparison has found *n_ab_* tags are shared by both repertoires. Pairwise type sharing (PTS) is then given by
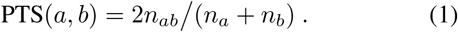

When *n_a_* and *n_b_* are nearly 60, the performance of PTS is excellent. For instance, when two parasites are completely different, *n_ab_* = 0, so PTS = 0; when two parasites are identical, and both repertoires have been fully sampled, *n_ab_* = *n_a_* = *n_b_*, so PTS = 1. However, when *n_a_* or *n_b_* is smaller (as is overwhelmingly the case in existing studies [7–13]) PTS is conservative and systematically underestimates the true overlap between repertoires [20]. For example, if we were able to fully sample two parasites that share 30 of their 60 *var* tags, their PTS would be 0.5. However, if *n_a_* = 37 and *n_b_* = 24 sequences are generated in repeated simulation (as in Fig. 1), the average resulting PTS is only 0.24, underestimating overlap by more than twofold. Furthermore, while intuitive, PTS is not underpinned by any statistical model, and therefore its uncertainty cannot be estimated. Outside the malaria literature, PTS is referred to as the Sørenson-Dice coefficient, having been independently published in the 1940s in the context of botany [21, 22], and is technically equivalent to the *F*_l_ score commonly used in evaluation of the effectiveness of prediction algorithms.

In this manuscript, I introduce a statistically rigorous alternative to PTS using Bayesian inference. By modeling the stochastic process by which repertoires are sampled, I show that this method produces unbiased *a posteriori* estimates of true repertoire overlap. I then show how the Bayesian framework can be used to estimate uncertainty and produce error bars which represent credible intervals, a Bayesian analog of confidence intervals. Finally, since each successful PCR amplification randomly samples just one of 60 available tags, I extend the Bayesian approach to compute the tradeoff between increasing PCR efforts and decreasing the uncertainty of repertoire overlap estimates. These calculations allow the cost of reagents and time to be weighed against scientific confidence, illustrating the use of this statistical framework for planning and budgeting experiments. Open-source code and a web tool are freely available (see Acknowledgements).

## II. METHODS

Suppose that there are two *P. falciparum* parasites, each with a repertoire of 60 *var* types. Our goal is to estimate the true repertoire overlap *s* (were we to fully sample each parasite) from the knowledge that *<u>n_a_</u>* samples from parasite *a* and *n_b_* samples from parasite *b* share *n_ab_* sequences. Due to the fact that the underlying sampling process is stochastic (Figure 1), our secondary goal is to quantify the uncertainty in the method’s estimates. Both goals can be met by writing down the process that creates the data in the first place. Therefore, in what follows, we will at first assume that the true overlap *s* is fixed, model the process of generating data via stochastic sampling, and use that model to compute a likelihood. We will then use Bayes’ Rule to compute the posterior probability for each value of *s*, given the evidence in the data and the likelihood computed in the first step.

Consider the following sampling process, written in the slightly more rigid and generic language of a probability textbook. Suppose that there are *s* special objects among a total of *N* objects. We draw *n* objects uniformly at random without replacement. The number of special objects chosen during this sampling procedure will be distributed according to a hypergeometric distribution, which we write as 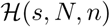.

First, with this definition in mind, consider drawing *n_a_ var* genes from parasite *a*’s 60 total. Of the 60 total, suppose that exactly *s* are considered special because they are also shared by parasite *b*.The number of shared sequences that are captured by sequencing parasite *a* will be a random variable 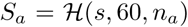. Depending on the luck of the draw, this number could be as small as zero, or as high as *s* or *n_a_* (whichever is smaller).

Now consider drawing *n_b_ var* genes from parasite *b*’s 60 total, in which exactly *s_a_* are special because they are shared by both parasites *and* were actually drawn during the sequencing of parasite *a*. This process is identical in construction to the process for sampling parasite *a*, but with *s_a_* special sequences instead of *s*, and so the number of shared sequences that are captured after sequencing both parasites will be 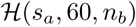. Substituting the random variable *S_a_* for a fixed value *s_a_*, which we derived in the paragraph above, yields a hypergeometric inside a hypergeometric, which means that the probability of a particular number of shared sequences in the samples *n_ab_* is given by these sequential (or nested) hypergeometric distributions,
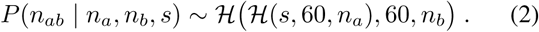

Reassuringly, one can switch the order in which the imagined sampling took place, first sequencing parasite *b* and then sequencing parasite *a*, or sequencing them both at once, and show that these are mathematically equivalent.

In practice, we want to go the other direction, and estimate *s* from our empirical measurements of *n_a_*, *n_b_*, and *n_ab_*. Since the distributions above allow us to compute the likelihood of empirical observations, given *s*, we use Bayes’ rule to formulate the posterior distribution for *s*,
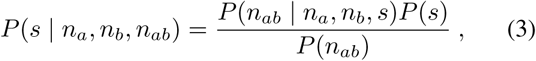

where *P*(*s*) is the prior distribution for overlap. In practice, we generally wish to remain agnostic about the level of overlap *s* and therefore we consider an uninformative prior *P*(*s*) ~ unif[0, 60], i.e. 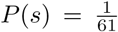. Using the law of total probability to rewrite the denominator, and canceling the factors of 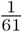, we get
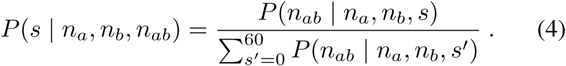

Each term on the right hand side of Eq. (4) can now be computed directly from the nested hypergeometric distributions in Eq. (2) as follows. To generate a specific empirical overlap *n_ab_*, two things must have happened in succession and independently of each other: first, *s_a_* of the original *s* shared sequences must have been sampled; and second, *n_ab_* of the intermediate *s_a_* shared sequences must then have been sampled. We therefore multiply these two hypergeometric probabilities. However, because this sequential process may occur for any value of the intermediate variable *s_a_*, we sum over all possible values of *s_a_*,
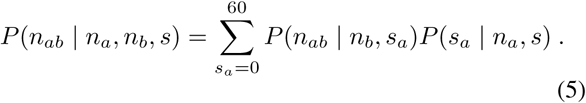

Thus, computing the posterior probability that the true overlap was *s*, given the empirical overlap between samples, is given by substituting Eq. (5) into Eq. (4), yielding
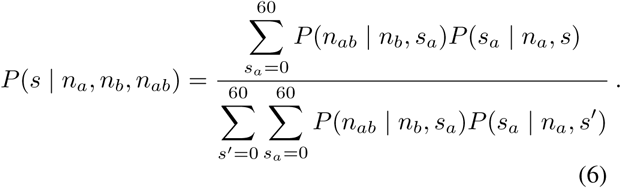

The term *P*(*s* | *n_a_*, *n_b_*, *n_ab_*) is a posterior distribution over s, meaning that it tells us the probability for each value of s, given the evidence provided by the actual data. While this equation appears notation-heavy, its inference requires only calls to the hypergeometric probability distribution. To illustrate this graphically, the posterior distribution is plotted for *n_a_* = 47, *n_b_* = 32, and *n_ab_* = 20 in Figure 2.

**FIG. 2.**
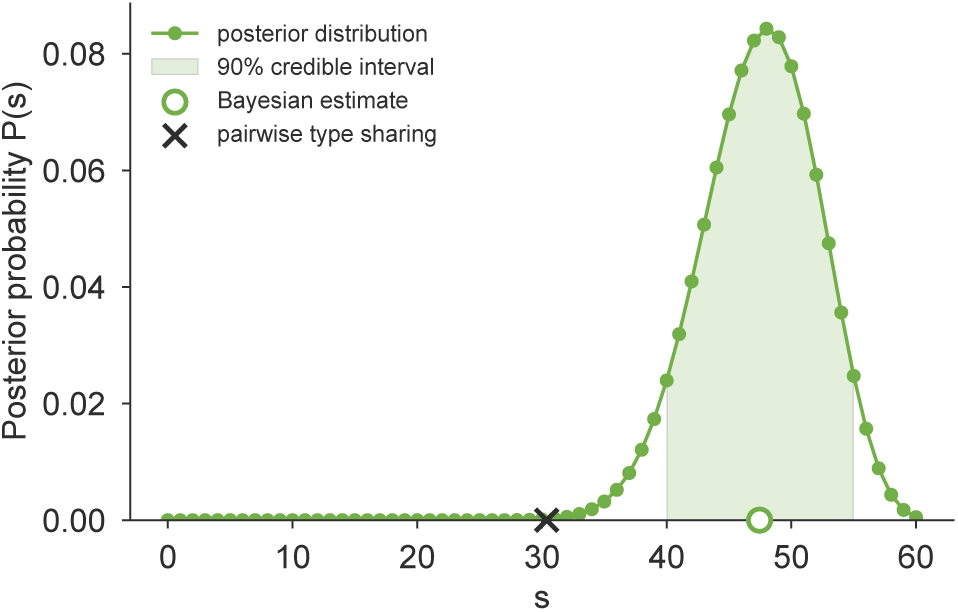
Inference and uncertainty using the posterior. The posterior distribution over *s* is plotted for the realistic scenario of *n_a_* = 47, *n_b_* = 32, and *n_ab_* = 20 [line; Eq. (6)]. The posterior mean provides our estimate of the true overlap 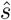 [open circle; Eq. (7)], and the interval accounting for at least 90% of the area under the posterior curve provides an equal-tailed 90% credible interval [shading; Eq. (8)]. The PTS estimate is shown for comparison [black cross; Eq. (1)], and is typically less than or equal to 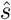.

The posterior distribution can now be used (i) to estimate the true value of *s*, and (ii) to quantify the uncertainty of that estimate. First, our estimate for the true value of *s*, which we call 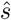, is the expected value of the posterior,
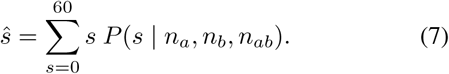

This value is typically (in 99.85% of all possible cases) larger than the estimate provided by PTS (Fig. 2).

The posterior distribution provides a convenient way to quantify the uncertainty associated with an estimate 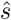. Intuitively, if the posterior is sharply peaked around 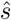, then our confidence in 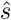 is high; if the posterior is broadly distributed then our confidence in 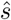 is low. Making use of the Bayesian construction once more, we compute a credible interval by finding the range of *s* values that account for 90% of the posterior probability (Fig. 2). Due to the fact that the posterior distribution is a discrete distribution over only 61 values, it is possible (indeed, highly probable) that no interval will contain exactly 90% of the probability. Nevertheless, we define a conservative equal-tailed 90% credible interval [*s*_min_, *s*_max_] as the smallest index *s*_min_ and the largest index *s*_max_ for which
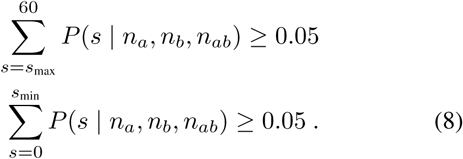

## III. RESULTS

### A. Estimator performance

We first demonstrate that the 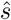 computed in Eq. (7) produces accurate estimates by simulating the sampling process with known *s* and evaluating our ability to accurately recover it. Specifically, for each simulation, we consider two *var* repertoires *a* and *b*, of 60 genes each, and specify a priori that they share exactly *s* sequences. We then choose the number of samples takes from each, *n_a_* and *n_b_* respectively, and draw from each repertoire uniformly at random, without replacement. These draws are compared to compute the number of empirically shared sequences *n_ab_*. Equation (7) is used to compute the Bayesian repertoire overlap (BRO) estimate 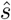, while Eq. (1) is used to compute PTS using the same data. These estimates are then compared to the true value of *s* to evaluate accuracy. Varying the values of *s*, *n_a_*, and *n_b_* allows us to quantify the performance of BRO and PTS in a variety of realistic sampling scenarios.

Figure 3 shows the results of this simulation for sampling rates of 30, 40, and 50 sequences, with two independent simulations at each value of *s*. Intuitively, both BRO and PTS become are more accurate when *n_a_* and *n_b_* are larger. However, the two methods’ behaviors are fundamentally different. When *n_a_* and *n_b_* are below 60, BRO provides estimates that are distributed around the true overlap, with variance increasing as sampling rates decrease. In contrast, PTS systematically underestimates the true overlap, while also showing increasing variance as sampling rates decrease [20]. For realistic sampling rates, BRO provides estimates centered at the true value, while PTS provides estimates centered below the true value.

**FIG. 3.**
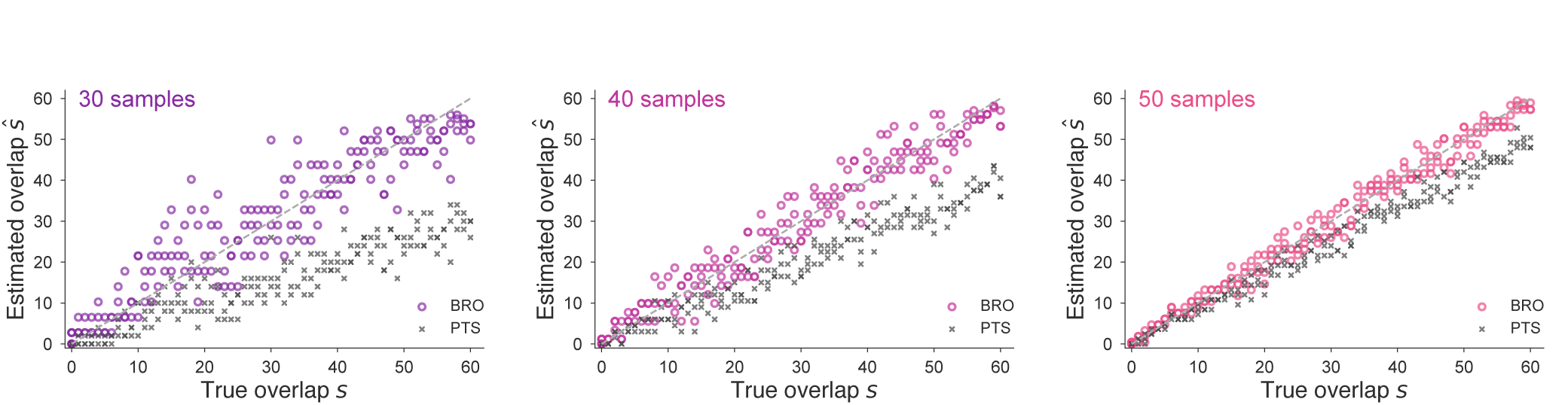
Bayesian repertoire overlap consistently estimates true overlap. Repertoires with true overlaps ranging from 0 to 60 were subsampled in simulations. As sampling rates increase from *n_a_* = *n_b_* = 30 (left) to 40 (middle) and to 50 (right), the estimates of BRO (colored circles) converge approximately symmetrically to the true values (dotted lines). Estimates from PTS (crosses) converge to the true values from below, systematically underestimating the true overlap. This bias is worse with lower sampling rates [20].

Credible intervals, which visually show uncertainty in each estimate, can also be easily computed from the simulations described above. For each simulation, Eq. (8) uses the posterior distribution over *s* to produce error bars around the point estimate 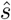, shown for sampling rates of 30, 40, and 50 in Figure 4. This illustrates the substantial reduction in uncertainty that comes with increased sampling rates. While all simulations shown here use *n_a_* = *n_b_*, this is by no means required, and in real data scenarios, is rare.

**FIG. 4.**
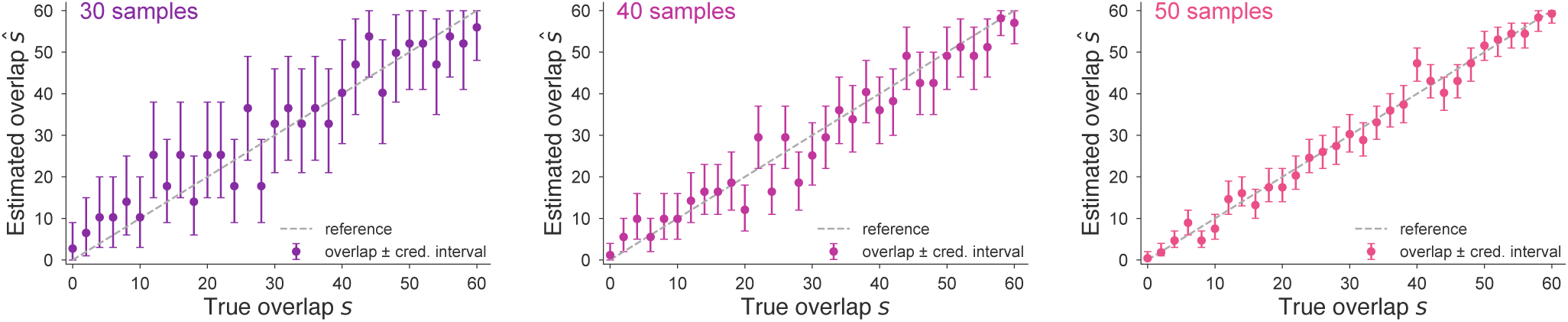
Credible intervals quantify uncertainty in overlap estimates. By using Eq. (8), 90% credible intervals are show above as error bars around the point estimates 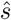 for varying true overlap *s*. As sampling rate increases from *n_a_* = *n_b_* = 30 (left) to 40 (middle) and to 50 (right), credible intervals shrink, indicating a reduction in uncertainty. In expectation, 90% of intervals cover the true overlap (dotted line).

### B. Revisiting past results

We now show how the methods of this paper can be used in practical contexts by applying them to data from three published studies. In particular, this reanalysis highlights the impact of variation in sampling rates across studies, which creates variable bias in PTS calculations and produces misleading results. However, we also show that while substitution of BRO for PTS sidesteps the bias problem, the ability to quantify uncertainty with error bars also highlights new problems. In short, the conclusions of previous studies may be worth reevaluating.

In 2007, Barry et al introduced PTS in an analysis of *var* data from Amele, Papua New Guinea [7]. In 2010, Albrecht et al included Barry’s data in a broader analysis of *var* data from Ariquemes, Brazil [8] which also included sequences from Kilifi, Kenya for comparison [17]. Each one of these studies, individually, sequenced parasite isolates to a particular target depth, yet the studies varied in their coverage of repertoires. Since the bias of PTS depends on the number of samples (Fig. 3; see also [20]), the variation of sampling rates across study populations means that different populations are biased downward by different amounts.

Albrecht et al conveniently provide *var* type data from all three studies, from which we can rebuild their first figure which shows a PTS comparison of five populations (Fig. 5; left). Overlaps between pairs of parasites can then be recomputed using BRO (Fig. 5; middle). The conclusions drawn from these two figures differ substantially.

**FIG. 5.**
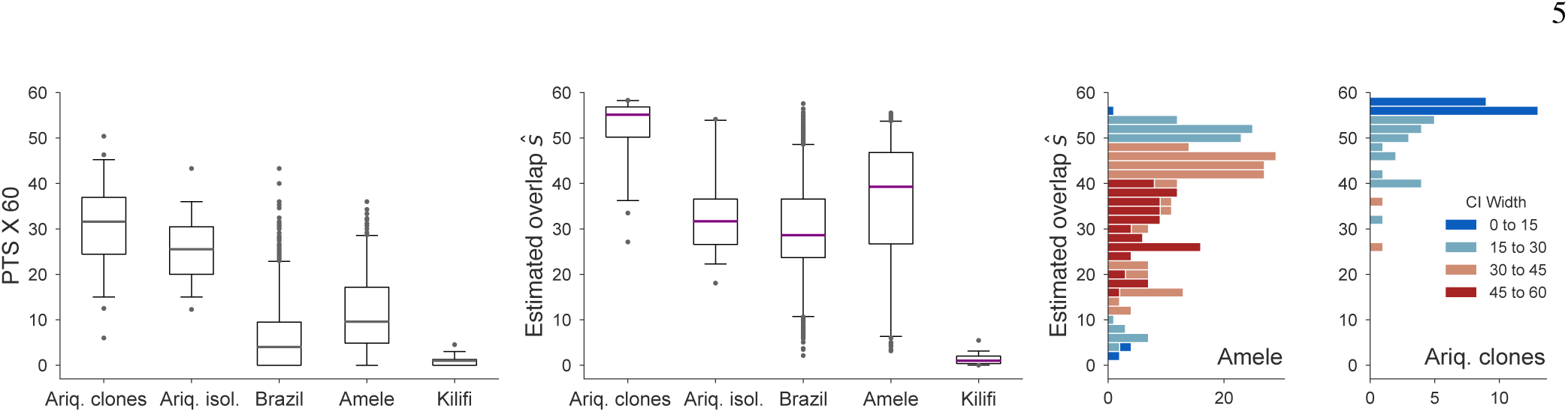
Reevaluation of published results. In 2010, Albrecht et al. compared *var* repertoires from 5 populations using pairwise type sharing (see Refs. [7, 8, 17] for original data details). (left) Reproduction of PTS analysis of [8], rescaled from [0,1] → [0, 60]. (middle) Reanalysis using Bayesian repertoire overlap Eq. (7). For all boxplots, boxes span inner quartiles; center lines show medians; whiskers extend to 2.5 and 97.5 percentiles. (right) Histograms of Bayesian repertoire overlap distributions from Amele and Ariquemes clones (data identical to those in middle boxplots) colored by width of credible interval Eq. (8), a measure of uncertainty. Differences in uncertainties are driven primarily by sampling rates: Amele samples average 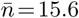 sequences per parasite while Ariquemes clones average 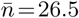.

First, according to PTS, identical clones from Ariquemes share only around 30 sequences with themselves, illustrating the downward bias produced by subsampling—clones ought to share all of their genes with their genetically identical siblings. Indeed, the reanalysis using BRO finds over 75% of overlap estimates to be greater than 50 (and over 50% over 55), far closer to what is expected.

Second, the inter-clone overlap and inter-isolate overlap distributions in Ariquemes appear to be similar and overlapping through the lens of PTS. However, the recalculation using BRO shifts the clones’ distribution dramatically upward but leaves the isolates’ distribution more or less untouched. This is due to the dramatic difference in *var* coverage: the average number of sequences per clone is 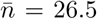 while for isolates it is 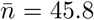, meaning that relatively different amounts of bias are inherited from PTS (illustrated in simulations in Fig. 3).

Finally, the distributions from Brazil (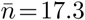) and Amele (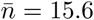) also shift dramatically upward when the bias of PTS is removed (Fig. 5; left, middle). However, this does not necessarily mean that they should be reinterpreted. For each pairwise comparison, Eq. (8) allows us to compute the width of the credible interval, *s*_max_ − *s*_min_ + 1, quantifying our uncertainty in each estimate. Due to low average coverage, the uncertainty of estimates in the Amele dataset tends to be extremely large (Fig. 5; right), with the majority of estimates showing an uncertainty greater than 30 sequences (50% overlap). For comparison, estimates from Thiès, Senegal [13] (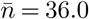) are also shown, whose dramatically lower uncertainty enables more confident conclusions to be drawn.

There are two main methodological findings that result from using rigorous and unbiased methods. First, the box-plots of Fig. 5 clearly illustrate that sampling rates can have a dramatic impact on findings, reinforcing the simulation results of Fig. 3. Second, uncertainty is an issue when 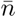 is too small, and datasets with low sampling rates may have such wide error bars that their estimates should not be trusted as shown in the histograms of Fig. 5, reinforcing the simulation results of Fig. 4. Additional sequencing efforts come at a cost, however, and so in the next subsection we use the methods of this paper to quantify the tradeoff between increased sequencing efforts and decreased uncertainty.

### C. The cost of reduced uncertainty

In the previous section, the reanalysis of published results shows clearly that the number of samples per parasite has a dramatic impact on the uncertainty (and therefore the interpretability) of painstakingly collected parasite sequence data. Naturally, increasing the sampling rates, *n_a_* and *n_b_*, decreases the uncertainty in 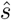, our estimate of *s* (Figure 4). However, additional samples cost time, effort, and money. Complicating matters, generating additional *var* sequences may or may not increase *n_a_*, since the previously sequenced *var* tags may be redundantly sequenced. Thus, there is a stochastic tradeoff between increased laboratory effort and decreased uncertainty about repertoire overlap, which we now calculate.

To obtain *var* tags, the DNA is PCR amplified using degenerate primers that are designed to capture all *var* genes with DBL*α* domains. This product is then cloned into a vector that allow single products to integrate, and these vectors are then transformed into bacteria and plated such that each colony contains one vector and one insert (see e.g. [13] for detailed methods). Therefore, among a large number of colonies, there are likely to be multiple colonies with the same *var* gene while some genes may not be covered by any colony. How many colonies should be separated and sequenced in order to get an accurate estimate of the repertoire overlap between two parasites? Put more formally, if we repeatedly perform an experiment in which we sequence *c* colonies each from two parasites and estimate their overlap 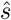, how much more accurate will 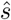 become if we increase *c*?

To answer this question, we split it into two parts. First, if we sequence *c* colonies, how many unique *var* genes *n* are we likely to have sampled? Second, what implications will this have for our repertoire overlap estimates, discussed in the previous section?

The first question can be answered by considering a process in which there are *k* = 60 distinct sequences in total and we draw *c* of them, one at a time, independently and with replacement. For a fixed *c*, we can compute the probability mass function for the number of distinct sequences by a straightforward recursion: At any point during the process of drawing sequences, if *n* distinct sequences have already been drawn, then the probability of drawing an already-discovered sequence is *n/k*, making the probability of drawing a new sequence 1 − *n/k*. Each draw is independent of the previous draws, so the incremental accumulation of distinct sequences can be written as a Markov chain with transition matrix *π* whose non-zero entries are
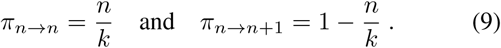

Initially, zero sequences have been drawn (*c* = 0), making *n* = 0 with probability 1. For each additional sequence drawn, the probability distribution over the number of distinct sequences evolves according to the transition matrix *π*, so that after *c* draws the distribution over distinct sequences is given by the entries of the vector x,
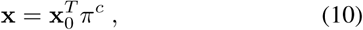

where x_0_ is initial condition vector of zeros, except for the entry corresponding to the state *n* = 0, which equals one. This allows us to analytically compute the distribution of the number of unique *var* genes sampled by a PCR process with *c* colonies. In other words, we now have a map between laboratory efforts *c* and the distribution of actual unique *var* genes sampled, and we write this as *P*(*n* | *c*). A variant of this problem was previously considered with the goal of computing the value of *c* that would cover at least 60% of each repertoire [23]. Although those calculations can be shown to produce incorrect estimates, Eq. (10) can be used to solve that problem variant as well. More widely, this general problem has been charmingly named *the coupon collector’s problem* by statisticians.

The second question focuses on the implications of Eq. (10), and specifically requires that we quantify how an increase in sequencing efforts *c* affects the noisy distribution of estimates 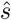. Intuitively, for low *c*, both *n_a_* and *n_b_* will tend to be small, leading to broad distributions of 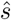 around the correct value of *s*. Similarly, as *c* grows very large, we expect the distribution of 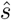 to concentrate on exactly *s*. This distribution, 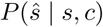, can be computed by integrating the distribution of estimates, conditioned on particular data, over the probability distribution of having produced those data, conditioned on *c* and *s*. Symbolically, the distribution of estimators 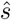, given true overlap *s* and colonies *c* is given by
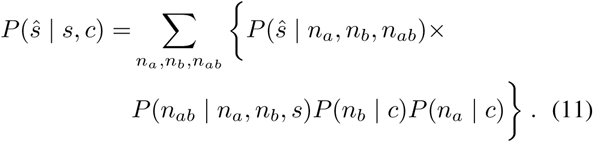

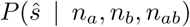 is the probability of getting a particular estimate 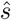, given information about coverage and overlap. In fact, this is a distribution concentrated at a single point, i.e., a Dirac *δ* function, since each triple (*n_a_*, *n_b_*, *n_ab_*) maps to exactly one point estimate 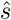. As a result, this term tells us the locations at which there will be probability mass, while the remaining terms in Eq. (11) tell us how much mass there will be at those locations. In other words, this distribution is a discrete probability distribution, and we have written down a fancy form of it above. By aggregating into bins, this distribution can be conveniently visualized as a histogram, which shows how the uncertainty of estimators depends on the true overlap *s* and the number of PCR colonies *c*. Figure 6 shows the effect of increasing sequencing efforts from a half plate (*c* = 48) to a full 96-well plate (*c* = 96) and beyond. These calculations succinctly quantify intuition: additional laboratory efforts lead to higher accuracy guarantees.

**FIG. 6.**
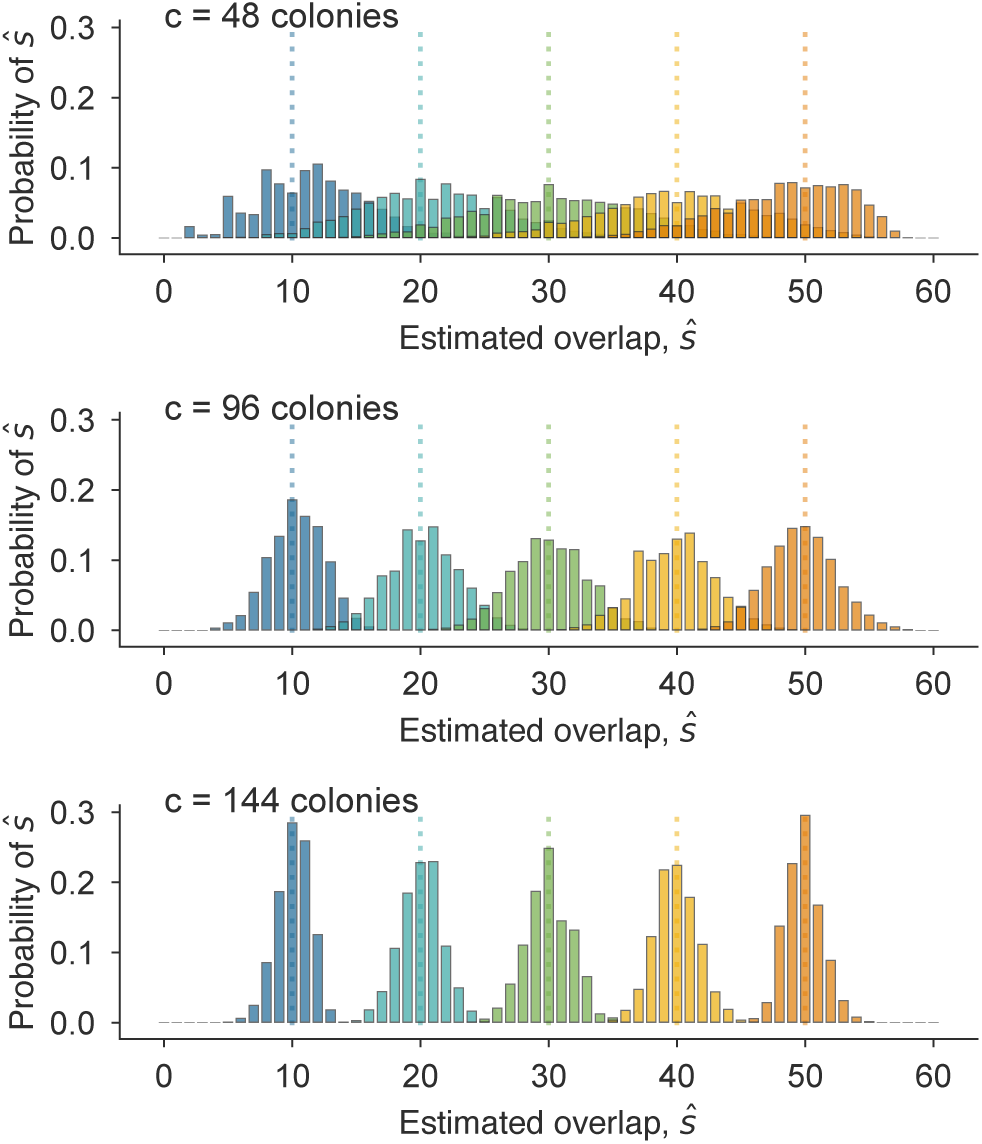
Quantifying the decrease in uncertainty from increased sequencing. Histograms show distributions of overlap estimates 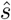, computed using Eq. (11), for various values of *s* which are indicated by color-matched dotted lines. While all estimates are distributed around the true values of s, increasing the number of colonies *c* from 48 (top) to 96 (middle) and to 144 (bottom) substantially decreases the error of estimates. For example the bottom plot shows that successfully sequencing *c* = 144 colonies from each parasite is guaranteed to produce estimates 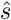 that are off by at most 5 in either direction of the true *s*.

The calculations and distributions in this section show how the Bayesian framework in this manuscript can also be used to plan sequencing studies and estimate study costs. If a desired downstream analysis of repertoire overlap requires results that are accurate to within a particular number of shared sequences, BRO methods can easily specify the sequencing efforts needed.

## IV. DISCUSSION

This manuscript places the estimation of repertoire overlap from imperfect samples on firm statistical ground. Past efforts used a convenient computation called pairwise type sharing (PTS), which was designed to be intuitive and conservative in its estimates of repertoire overlap. Here, we clearly define a stochastic process that generates the data we observe, opening the door to more rigorous Bayesian inference. In particular, Eq. (7) provides point estimates of true repertoire overlap, while Eq. (8) provides error bars and uncertainty estimates via credible intervals. Figures 3 and 4 show the consistency and accuracy of these calculations across simulated sampling regimes in which the correct answer is known.

Bayesian repertoire overlap is also useful in real-data scenarios, when the correct answer is unknown. By revisiting previously published studies [7, 8, 13, 17] we showed that the conclusions that can confidently be drawn may change (Fig. 5 left, middle) or disappear (Fig. 5 right). In particular, the reanalysis points to a clear recommendation for the design of future studies: the number of unique sequences per isolate should be at least 30. Since each additional PCR product may not contribute an additional unique sequence, we again used the Bayesian framework to translate increased PCR efforts to decreased uncertainty (Fig. 6). Accuracy requirements can now be weighed against laboratory costs during the planning of studies.

While BRO clearly outperforms PTS in practical contexts, it is also more cumbersome to compute. Indeed, PTS can be calculated on the back of an envelope while Eq. (7) requires a computer, or at least a lot more envelopes. However, as it turns out, there are only around 77,500 possible combinations of *n_a_*, *n_b_*, and *n_ab_*, which means that a lookup table of every conceivable 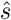 value can be computed on a laptop in minutes and attached to an email. Links to open-source code and a convenient web tool can be found in the Acknowledgements.

The models introduced in this paper are as correct as their assumptions, which we now revisit. During the construction of the Bayesian repertoire overlap, we assumed that our prior distribution *P*(*s*) was uniform, meaning that we treated each possible level of overlap as equally likely. This is easily defensible in practice, as any other choice would introduce unacceptable bias. However, in computing the tradeoff between sequencing effort and uncertainty, we also assumed that each sequence in each repertoire was just as likely to have been sampled, which may or may not be true. Due to the fact that sequencing takes place via PCR using degenerate primers, the effects of primer bias may cause some sequences to be amplified more often than others. Fully addressing this possibility would require that we modify the probabilities in both the coupon collector’s problem and the repertoire subsampling processes. In fact, it is possible that deviations from our modeling assumptions (i.e., primer bias) could be inferred from the data themselves. This is an interesting topic, but remains outside the scope of the current paper.

The calculations in this paper also assume a *var* repertoire size of 60. In reality, this assumption is routinely violated. Simulation studies (not shown) suggest that small fluctuations in the total assumed repertoire size make little practical difference. Nevertheless, as larger whole-genome datasets become available, an additional prior over the distribution of repertoire sizes could improve estimates further.

More practically, the assumption that the *var* repertoire size is 60 makes the methods of this paper useless in the context of complex infections with multiple parasite genomes [12, 17]. In cases where the multiplicity of infection is known, overlap estimates could be computed using generalizations of the statistics in this paper, computing overlap between infections (instead of between parasites). However, this would be complicated by possible overlap of parasite repertoires within each infection. Nevertheless, development of such methods would be especially useful in the context of *var*-based epidemiological studies.

Finally, it is possible that unbiased methods for comparing *var* cDNA could be derived from the framework presented here. In contrast to uniform sampling of gDNA, non-uniform results are precisely the point of sequencing *var* cDNA, in order to link differential *var* expression patterns with clinical phenotypes. This would require jointly estimating *var* expression *and* repertoires, but in studies with both cDNA and gDNA (e.g. [13]) such calculations may be feasible.

## V. ACKNOWLEDGEMENTS

DBL was supported by the Ruth and Sidney Weiss Fund, and would like to thank Amy K. Bei, Samuel F. Way, and Caroline O. Buckee for helpful conversations and suggestions. DBL also warmly thanks the authors of Refs. [7, 8] whose commitment to open science and methods means that their data were freely available for re-analysis.

Open-source code is freely available in Python at <u>github.com/dblarremore/BayesianRepertoireOverlap</u>. A web tool version produces estimates, credible intervals, and figures like Fig. 2 and is available at <u>bro.colorado.edu</u>.

